# Energetic Demands Regulate Sleep-Wake Rhythm Circuit Development

**DOI:** 10.1101/2023.09.19.558472

**Authors:** Amy R. Poe, Lucy Zhu, Si Hao Tang, Ella Valencia, Matthew S. Kayser

## Abstract

Sleep and feeding patterns lack strong daily rhythms during early life. As diurnal animals mature, feeding is consolidated to the day and sleep to the night. In *Drosophila*, circadian sleep patterns are initiated with formation of a circuit connecting the central clock to arousal output neurons; emergence of circadian sleep also enables long-term memory (LTM). However, the cues that trigger the development of this clock-arousal circuit are unknown. Here, we identify a role for nutritional status in driving sleep-wake rhythm development in *Drosophila* larvae. We find that in the 2^nd^ instar larval period (L2), sleep and feeding are spread across the day; these behaviors become organized into daily patterns by the 3^rd^ instar larval stage (L3). Forcing mature (L3) animals to adopt immature (L2) feeding strategies disrupts sleep-wake rhythms and the ability to exhibit LTM. In addition, the development of the clock (DN1a)-arousal (Dh44) circuit itself is influenced by the larval nutritional environment. Finally, we demonstrate that larval arousal Dh44 neurons act through glucose metabolic genes to drive onset of daily sleep-wake rhythms. Together, our data suggest that changes to energetic demands in developing organisms trigger the formation of sleep-circadian circuits and behaviors.

## Introduction

The development of behavioral rhythms such as sleep-wake patterns is critical for brain development^1^. Indeed, early life circadian disruptions in rodents negatively impacts adult behaviors, neuronal morphology, and circadian physiology^2–4^. Likewise, in humans, disruptions in sleep and rhythms during development are a common co-morbidity in neurodevelopmental disorders including ADHD and autism^5–8^. Although mechanisms encoding the molecular clock are well understood, little is known about how rhythmic behaviors first emerge^1,9–11^. In particular, cues that trigger the consolidation of sleep and waking behaviors as development proceeds are unclear^12–15^.

A key potential factor in the maturation of sleep patterns is the coincident change in feeding and metabolism during development. Early in development, most young animals must obtain enough nutrients to ensure proper growth^16^. Yet, developing organisms must also sleep to support nervous system development^1^. These conflicting needs (feeding vs. quiescence) result in rapid transitions between sleeping and feeding states early in life^17,18^. As development proceeds, nutritional intake and storage capacity increase, allowing for the consolidation of feeding and sleep to specific times of day^19^. These changes in nutritional storage capacity are likely conserved as mammalian body composition and nutritional capacity change over infant development^20^ and *Drosophila* show rapid increases in overall larval body size including the size of the fat body (used for nutrient storage) across development^21,22^. However, the role that developmental change in metabolic drive plays in regulating the consolidation of behavioral rhythms is not known.

In adult *Drosophila*, sleep and feeding behaviors are consolidated to specific times of day with flies eating more in the day than the night^23^. Yet, early in development, sleep in 2^nd^ instar *Drosophila* larvae (L2) lacks a circadian pattern^24^. We previously determined that sleep-wake rhythms are initiated in early 3^rd^ instar *Drosophila* larvae (L3) (72 hr AEL). DN1a clock neurons anatomically and functionally connect to Dh44 arousal output neurons to drive the consolidation of sleep in L3. Development of this circuit promotes deeper sleep in L3 resulting in the emergence of long-term memory (LTM) capabilities at the L3 stage but not before^24^.

Here, we identify the cues that trigger the emergence of the DN1a-Dh44 circuit and the consolidation of sleep-wake rhythms in *Drosophila* larvae. We demonstrate that developmental changes to energetic demands drive consolidation of periods of sleep and feeding across the day as animals mature. While endogenous deeper sleep in L3 facilitates LTM, we find that experimentally inducing deep sleep prematurely in L2 is detrimental to development and does not improve LTM performance. Additionally, we demonstrate that DN1a-Dh44 circuit formation is developmentally plastic, as rearing on an insufficient nutritional environment prevents establishment of this neural connection. Finally, we find that Dh44 neurons require glucose metabolic genes to promote sleep-wake rhythm development, suggesting that these neurons sense the nutritional environment to promote circadian-sleep crosstalk.

## Results

### Energetic demands limit developmental onset of rhythmic behaviors

The emergence of circadian sleep-wake rhythms in *Drosophila* larvae is advantageous as it enables long-term memory capabilities at the L3 stage^24^. Less mature larvae (L2) do not exhibit consolidated sleep-wake patterns, prompting us to ask why rhythmic behaviors do not emerge earlier in life. To determine if the absence of rhythmic sleep-wake patterns in L2 might be related to feeding patterns, we examined larval feeding rate (# mouth hook contractions in a 5-minute period) under constant conditions at 4 times across the day (Circadian Time [CT] 1, CT7, CT13, and CT19) in developmentally age-matched 2^nd^ (L2) and early 3^rd^ instar (L3; 72 hr AEL) larvae. While we observed no differences in feeding rate across the day in L2, we found that L3 show diurnal differences in feeding with higher feeding rate during the subjective day compared to the subjective night (Figures 1A and 1B). Analysis of total food intake indicated that L2 consume the same amount at CT1 and CT13; however, L3 consume more at CT1 than at CT13 (Figure S1A). To assess whether the daily pattern of feeding in early L3 is dependent on the canonical circadian biological clock, we examined feeding rate in null mutants for the clock gene *tim*^25^. We observed no differences in feeding rate across the day in L3 *tim* mutants indicating that the daily feeding pattern requires a functioning molecular clock (Figure 1C). These findings underscore the tight relationship between sleep and feeding across development as diurnal differences in sleep emerge concurrently at the L3 stage^24^.

**Figure 1:**
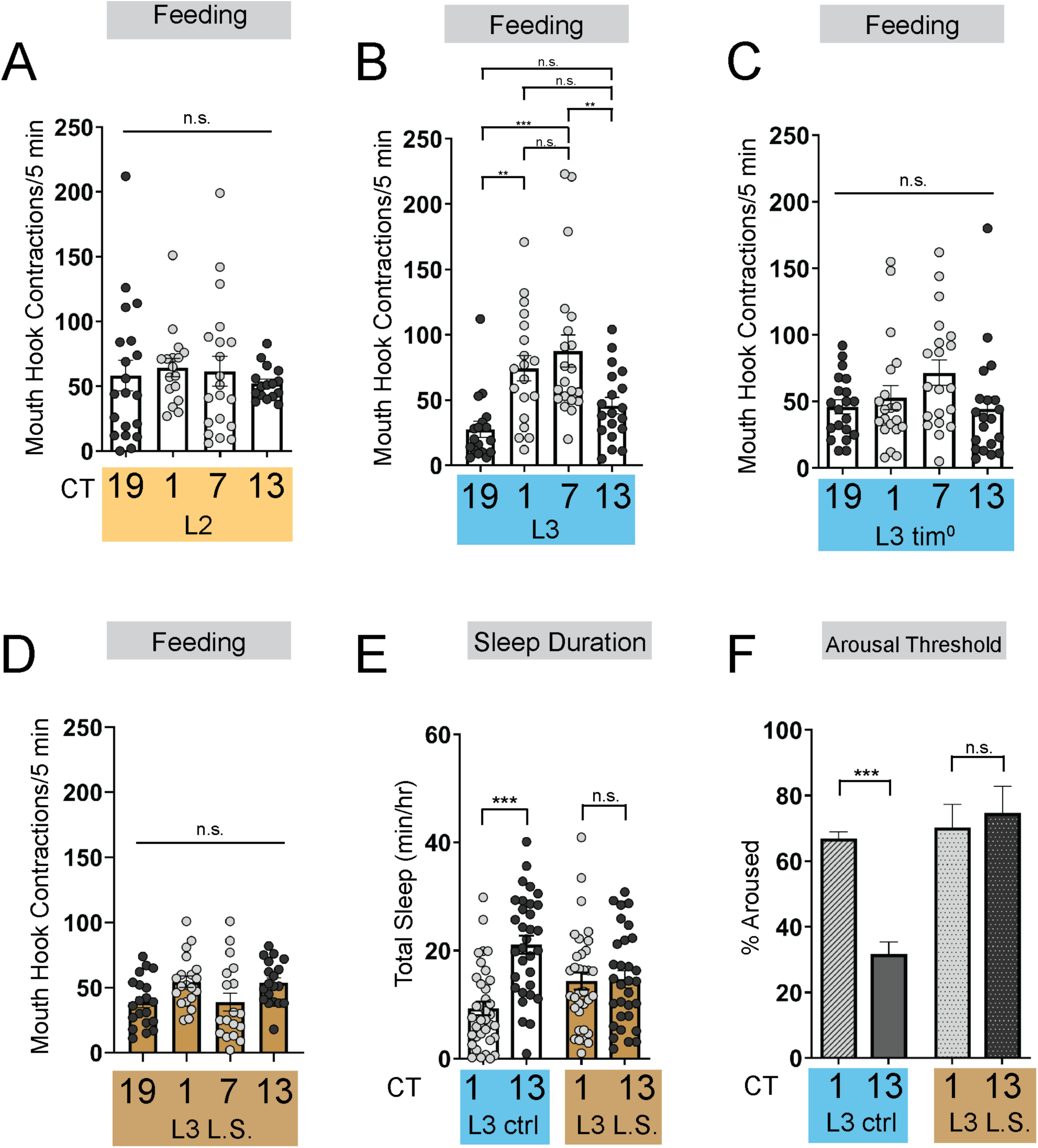
Energetic drive limits sleep rhythm development. **(A-D)** Feeding rate (# of mouth hook contractions per 5 min) of L2 controls (A), L3 raised on regular (ctrl) food (B), L3 clock mutants (C), and L3 raised on low sugar (L.S.) food (D) across the day. **(E)** Sleep duration at CT1 and CT13 in L3 raised on regular (ctrl) and L.S. food. **(F)** Arousal threshold at CT1 and CT13 in L3 raised on regular (ctrl) and L.S. food. A-D, n=18-20 larvae; E, n=29-34 larvae; F, n=100-172 sleep episodes, 18 larvae per genotype. One-way ANOVAs followed by Sidak’s multiple comparisons tests [(A-D)]; Two-way ANOVAs followed by Sidak’s multiple comparison test [(E-F)]. For this and all other figures unless otherwise specified, data are presented as mean ± SEM; n.s., not significant, \**P*<0.05, \*\**P*<0.01, \*\*\**P*<0.001.

To further investigate how the emergence of circadian sleep is related to changes in feeding patterns during development, we asked whether enforcing a constant (immature) feeding pattern at the L3 stage affects sleep-wake rhythms. First, we devised a nutritional paradigm with reduced sugar content but otherwise normal food (low sugar, 1.2% glucose, L.S.). Critically, this paradigm did not affect any measures of larval growth or development (Figures S1D-S1F) in contrast to numerous other diets that were assessed. We found that the feeding pattern in L3 raised on L.S. food closely resembled that of L2 on normal food (8% glucose), with feeding spread out across the day (Figure 1D) (Figure S1A). Compared to L3 raised on control food, this paradigm was also associated with loss of diurnal differences in sleep duration, sleep bout number, and arousal threshold (indicating less deep sleep) (Figures 1E and 1F) (Figures S1B and S1C). Next, to avoid chronic effects of this dietary manipulation, we acutely stimulated feeding in L3 reared on normal food using thermogenetic activation of NPF+ neurons (Figure 2A; Figure S2A)^26,27^. Like L.S. conditions, enforcing a constant feeding pattern through NPF+ neuron activation led to loss of sleep rhythms and loss of deep sleep (higher arousal threshold) in L3 (Figures 2B-2E; Figures S2B-S2E).

**Figure 2:**
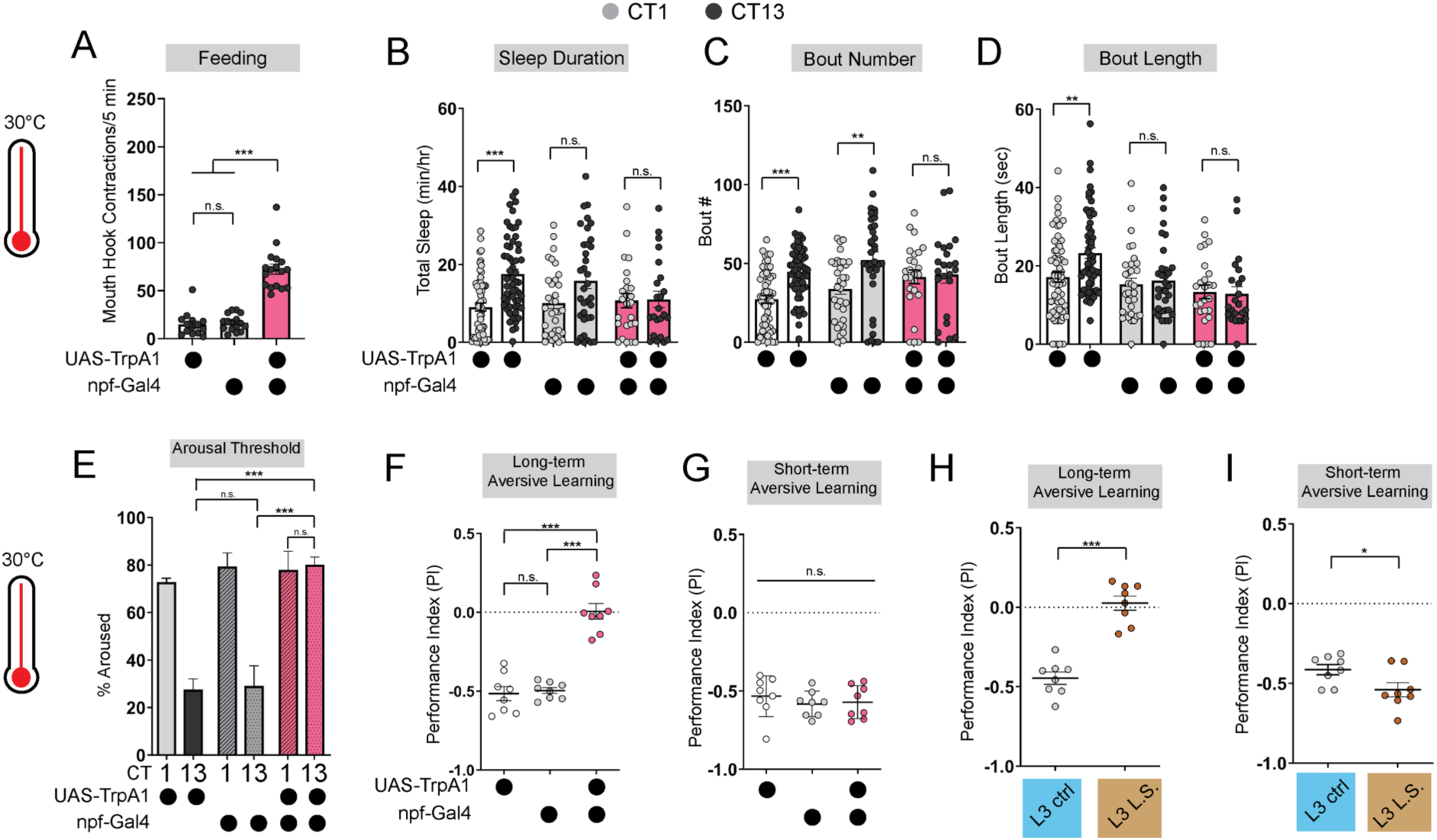
Immature feeding strategies limit LTM. **(A)** Feeding rate of L3 expressing *npf-*Gal4>*UAS-TrpA1* and genetic controls at 30°C at CT13. **(B-D)** Sleep duration (B), bout number (C), and bout length (D) of L3 expressing *npf-*Gal4>*UAS-TrpA1* and genetic controls at 30°C. **(E-G)** Arousal threshold (E), long-term aversive memory performance (F), and short-term aversive memory performance (G) in L3 expressing *npf-*Gal4>*UAS-TrpA1* and genetic controls at 30°C. **(H,I)** Short-and long-term term aversive memory performance in L3 raised on ctrl and L.S. food. A, n=18-20 larvae; B-D, n=24-61 larvae; E, n=125-160 sleep episodes, 18 larvae per genotype; F-I, n=8 PIs (240 larvae) per genotype. One-way ANOVAs followed by Sidak’s multiple comparisons tests [(A) and (E-G)]; Two-way ANOVAs followed by Sidak’s multiple comparison test [(B-D)]; unpaired two-tailed Student’s *t*-test [(H) and (I)].

Disruption to circadian sleep and/or to deeper sleep stages during development is associated with impairments in long-term memory formation^4,28–31^. We next asked if the loss of deeper sleep observed in animals adopting an immature (constant) feeding strategy through either the dietary paradigm or NPF+ neuron activation affects long-term memory (LTM). Consistent with deeper sleep stages being necessary for LTM, we observed a loss of LTM in either the NPF+ neuron activation (Figure 2F; Figures S2F-2I) or under L.S. conditions (Figure 2H; Figures S1G-S1I); short-term memory (STM) was intact in both manipulations (Figures 2G and 2I). These findings suggest that immature feeding strategies preclude the emergence of sleep rhythms and LTM. Together, our data indicate that consolidated periods of sleep and feeding emerge due to developmentally dynamic changes in energetic demands.

### Deeper sleep stages are energetically disadvantageous in L2

To investigate if promoting deep sleep at night in L2 can enable precocious LTM abilities, we fed L2 larvae the GABA-A agonist gaboxadol^32,33^. Gaboxadol feeding induced deeper sleep in L2 (as reflected by an increase in arousal threshold) although sleep duration was unchanged. Despite achieving deeper sleep, LTM was still not evident in L2 (Figures 3A,B,E; Figures S3A and S3B); however, in contrast to L3, L2 on gaboxadol failed to develop normally (Figures 3C and 3D). Next, to avoid chronic pharmacological manipulations altogether, we acutely stimulated sleep-inducing neurons using thermogenetic approaches. We found that acute activation of these neurons caused an increase in sleep duration and bout length with a decrease in arousal threshold (Figures 3F and 3G; Figures S3C-S3H). However, inducing deeper sleep in L2 via genetic approaches likewise did not improve LTM performance (Figure 3H; Figures S3I-S3K) despite STM being intact (Figure S3L). Moreover, as with gaboxadol feeding in L2, thermogenetic induction of deeper sleep stages disrupted larval development (Figures 3I and 3J). These data suggest that sleep cannot be leveraged to enhance cognitive function prematurely because prolonged periods of deep sleep are not energetically sustainable at this stage.

**Figure 3:**
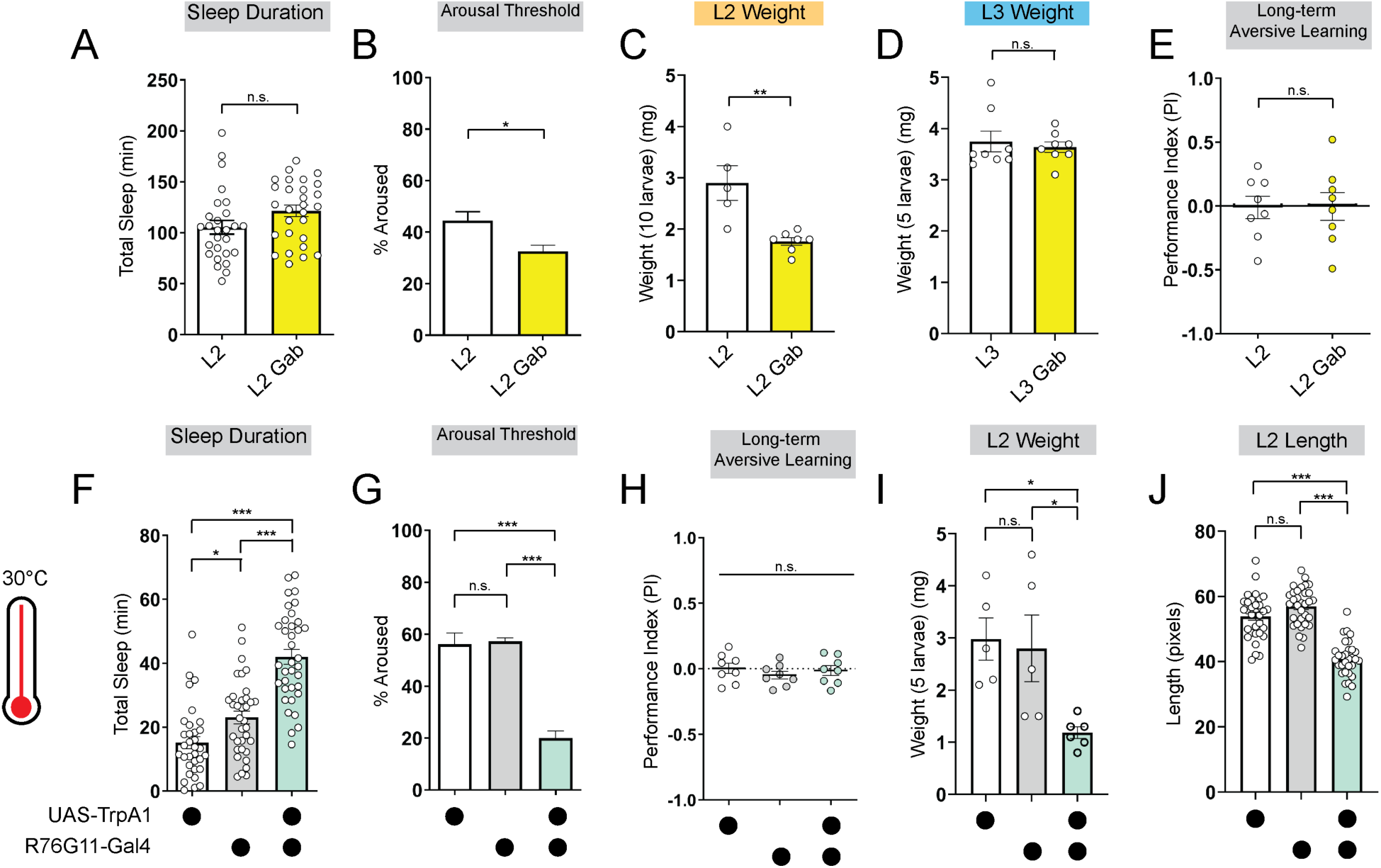
Deeper sleep in L2 is energetically disadvantageous. **(A, B)** Sleep duration (A) and arousal threshold (B) of L2 control fed vehicle control (L2) or Gaboxadol (L2 Gab). **(C, D)** Total body weight of L2 (C) (in groups of 10) or L3 (D) (in groups of 5) fed vehicle control or Gaboxadol (Gab). **(E)** Long-term aversive memory performance in L2 fed vehicle control (L2) or Gaboxadol (L2 Gab). **(F, G)** Sleep duration (F) and arousal threshold (G) of L2 expressing *R76G11-*Gal4>*UAS-TrpA1* and genetic controls at 30°C. **(H)** Long-term aversive memory performance of L2 expressing *R76G11-*Gal4>*UAS-TrpA1* and genetic controls at 30°C. **(I, J)** Total body weight (I) and total body length (J) of L2 expressing *R76G11-*Gal4>*UAS-TrpA1* and genetic controls at 30°C. A, n=28 larvae; B, n=110-220 sleep episodes, 18 larvae per genotype; C, n=5-7 groups (50-70 larvae); D, n=8 groups (40 larvae); E, n=8 PIs (240 larvae) per genotype; F, n=33-36 larvae; G, n=234-404 sleep episodes, 30-40 larvae per genotype; H, n=8 PIs (240 larvae) per genotype; I, n=5 groups (25 larvae); J, n=31-32 larvae. Unpaired two-tailed Student’s *t*-tests [(A-E)]; one-way ANOVAs followed by Sidak’s multiple comparisons tests [(F-J)].

### DN1a-Dh44 circuit formation is developmentally plastic

We previously determined that Dh44 arousal neurons anatomically and functionally connect to DN1a clock neurons at the L3 stage^24^. Therefore, we examined the functional connectivity between clock and arousal loci in the setting of nutritional perturbation by expressing ATP-gated P2X2 receptors^34^ in DN1a neurons and GCaMP6 in Dh44 neurons. As expected, activation of DN1as in L3 raised on control food caused an increase in calcium in Dh44 neurons (Figures 4A and 4B)^24^. In contrast, in L3 raised on L.S. conditions, activation of DN1as no longer elicited a response in Dh44 neurons (Figures 4C and 4D). Thus, the nascent connection underlying circadian sleep in *Drosophila* is developmentally plastic: in an insufficient nutritional environment, this connection is not functional, presumably to facilitate a more constant feeding pattern that fulfills energetic needs of the animal. However, this feeding pattern eschews deep sleep at the expense of LTM.

**Figure 4:**
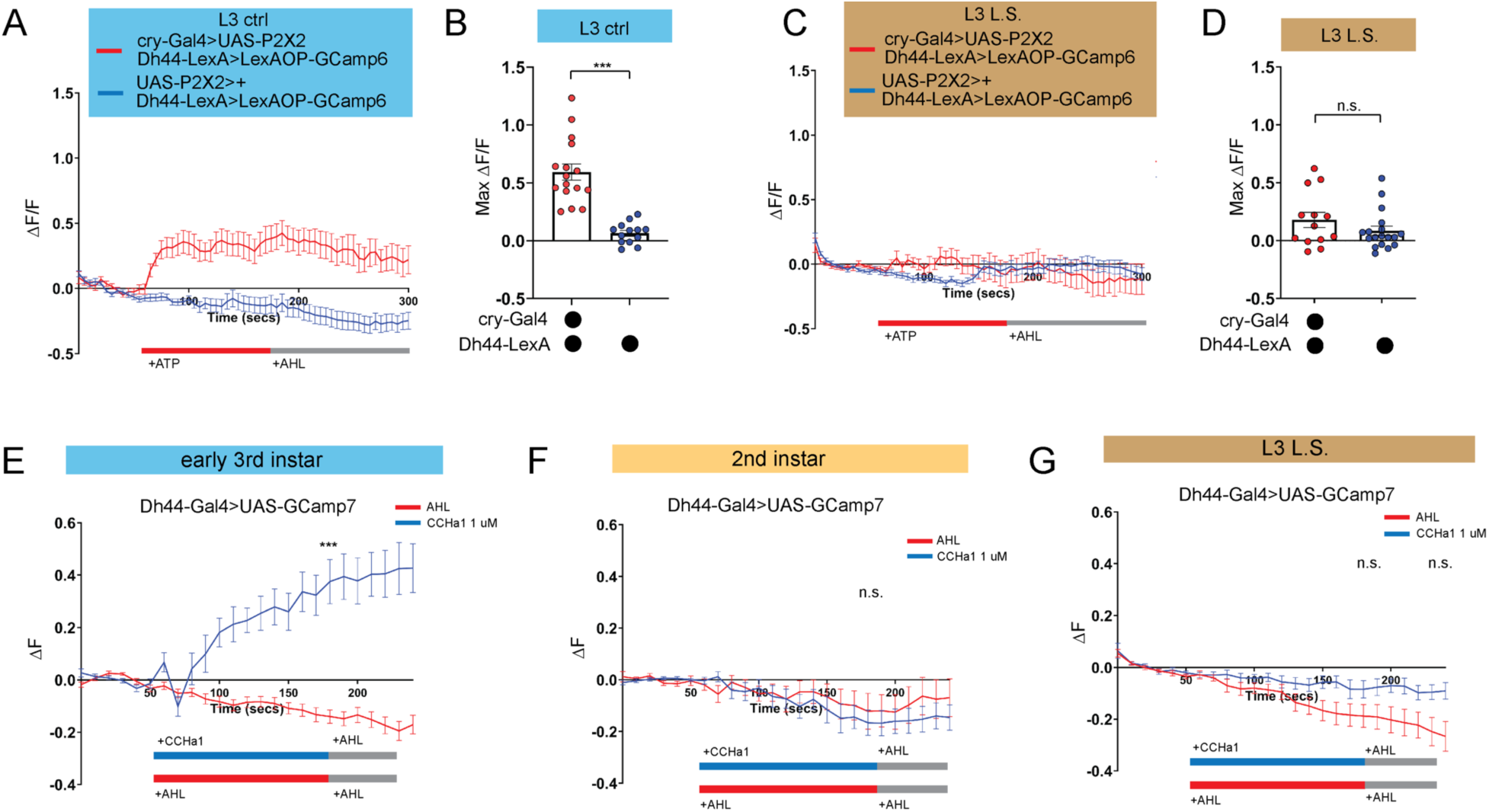
DN1a-Dh44 circuit formation is developmentally plastic. **(A, C)** GCaMP6 signal in Dh44 neurons with activation of DN1a neurons in L3 controls (A) and L3 raised on L.S. food (C). Red bar indicates ATP application and gray bar indicates AHL application. **(B, D)** Maximum GCaMP change (τιF/F) for individual cells in L3 controls (B) and L3 raised on L.S. food (D). **(E-G)** GCaMP7 signal in Dh44 neurons during bath application of 1 µM CCHa1 synthetic peptide in L3 controls (E), L2 controls (F) and L3 raised on L.S. food (G) brains. Red/blue bar indicates timing of CCHa1 (blue) or buffer (AHL, red) application and gray bar indicates timing of washout AHL application. A-D, n=12-18 cells, 8-10 brains; E-G, n=11-15 cells, 5-10 brains. Unpaired two-tailed Student’s *t*-tests [(B) and (D)]; Mann-Whitney U test [(E-G)].

We previously determined that release of *CCHamide-1* (*CCHa1*) from DN1as to *CCHa1mide-1 receptor* (*CCHa1-R*) on Dh44 neurons is necessary for sleep-wake rhythms in L3^24^. Our data support a model in which the downstream Dh44 neurons are poised to receive clock (DN1a) input as soon the connection forms between these cellular populations. To test this idea directly, we next asked whether Dh44 neurons in L2, before the DN1a-Dh44 connection has formed, are competent to receive the CCHa1 signal. CCHamide-1 peptide was bath applied onto dissected larval brains expressing *UAS-GCaMP7* in Dh44 neurons. As anticipated, we observed an increase in intracellular calcium in early L3 (Figure 4E); surprisingly, CCHa1 application did not alter calcium levels in Dh44 neurons in L2 (Figure 4F), indicating that Dh44 neurons are not capable of receiving CCHa1 input prior to early L3. Moreover, CCHa1 application in L3 reared on L.S. also failed to elicit a response in Dh44 neurons (Figure 4G) suggesting that sub-optimal nutritional milieu influences the development of Dh44 neuronal competency to receive clock-driven cues. Thus, our data indicate that the nutritional environment influences DN1a-Dh44 circuit development.

### Dh44 neurons require glucose metabolic genes to regulate sleep-wake rhythms

How do developing larvae detect changes in their nutritional environments to drive the circadian consolidation of sleep and feeding at the L3 stage? In adult *Drosophila*, Dh44 neurons act as nutrient sensors to regulate food consumption and starvation-induced sleep suppression through the activity of both glucose and amino acid sensing genes^35–38^. Indeed, Dh44 neurons themselves are activated by changes in bath application of nutritive sugars^35^. We next asked whether Dh44 neurons in L3 require metabolic genes to regulate sleep-wake rhythms. In L3 raised on regular food, we conducted an RNAi-based candidate screen of different glucose and amino acid sensing genes known to act in adult Dh44 neurons. Sleep duration at CT1 and CT13 was assessed with knockdown of glucose metabolic genes (*Hexokinase-C*, *Glucose transporter 1,* and *Pyruvate kinase*) or amino acid sensing genes (*Gcn2* and its downstream target ATF4 or *cryptocephal*) in Dh44 neurons. We found that knockdown of glucose metabolism genes, *Glut1*, *Hex-C* and *PyK*, in Dh44 neurons resulted in a loss of rhythmic changes in sleep duration and bout number (Figures 5A-5C; Figures S4A-S4F) with no effect on L3 feeding in the *Hex-C* manipulation (Figure 5D). Additionally, knockdown of *Hex-C* in DN1as did not disrupt rhythmic changes in sleep duration in L3 (Figures S4G-I) suggesting a specialized role for glucose metabolism in Dh44 neurons for sleep-wake rhythm maturation. Manipulation of glucose metabolic genes in L2 did not affect sleep duration at CT1 and CT13 (Figures 5E-5G; Figures S4J-S4O) providing evidence that nutrient sensing is not required at this stage to regulate sleep. In contrast to their role in adult Dh44 neurons, knockdown of amino acid sensing genes, *Gcn2* and *crc*, in Dh44 neurons did not disrupt rhythmic changes in sleep duration and bout number in L3 (Figures 5H and 5I; Figures S5A-S5D) suggesting that Dh44 neurons may not act through amino acid sensing pathways to regulate sleep-wake rhythm development. Thus, Dh44 neurons require glucose metabolic genes to drive sleep-wake rhythm development. Our data indicate that the emergence of daily sleep-wake patterns is regulated by developmental changes in energetic capacity, and suggest that Dh44 neurons may be necessary for sensing of larval nutritional environments.

**Figure 5:**
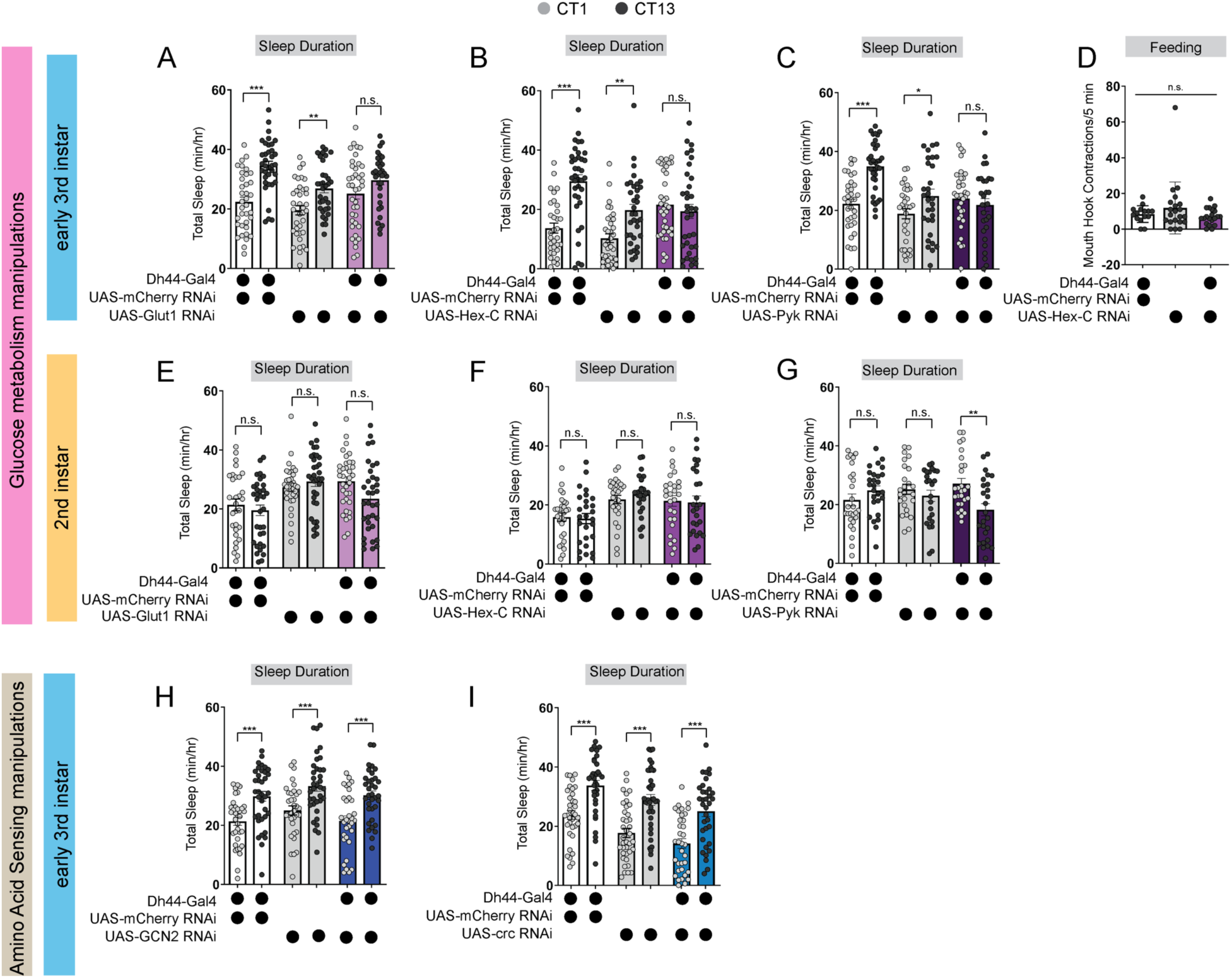
Dh44 neurons require glucose metabolic genes to regulate sleep-wake rhythms. **(A-C)** Sleep duration in L3 expressing *UAS-Glut1-RNAi* (A), *UAS-Hex-C-RNAi* (B), and *UAS-Pyk-RNAi* (C) with *Dh44-Gal4* and genetic controls at CT1 and CT13. **(D)** Feeding rate (# of mouth hook contractions per 5 min) of L3 expressing *UAS-Hex-C-RNAi* with *Dh44-Gal4* and genetic controls at CT13. **(E-G)** Sleep duration in L2 expressing *UAS-Glut1-RNAi* (E), *UAS-Hex-C-RNAi* (F), and *UAS-Pyk-RNAi* (G) with *Dh44-Gal4* and genetic controls at CT1 and CT13. **(H, I)** Sleep duration in L3 expressing *UAS-GCN2-RNAi* (H) and *UAS-crc-RNAi* (I) with *Dh44-Gal4* and genetic controls at CT1 and CT13. A-C, n=32-40 larvae; D, n=20 larvae; E-I, n=32-40 larvae. Two-way ANOVAs followed by Sidak’s multiple comparison test [(A-C) and (E-I)]; One-way ANOVAs followed by Sidak’s multiple comparisons tests [(D)].

## Discussion

Nutritional environment and energetic status exert profound effects on sleep patterns during development, but mechanisms coupling sleep to these factors remain undefined. We report that the development of sleep-circadian circuits depends on organisms achieving sufficient nutritional status to support the consolidation of deep sleep at night (Figure S6). Our data demonstrate that larval Dh44 neurons require glucose metabolic genes but not amino acid sensing genes to modulate sleep-wake rhythms. Larval Dh44 neurons may therefore have distinct functions from their adult counterparts for integrating information about the nutritional environment through the direct sensing of glucose levels to modulate sleep-wake rhythm development. Maintaining energy homeostasis and sensing of the nutritional environment are likely conserved regulators of sleep-wake rhythm development, as young mice exposed to a maternal low-protein diet show disruptions in night-time sleep architecture and energy expenditure later in life^39,40^.

Together with our previously published work, our findings support a model in which changes in both overall circuit development and molecular changes in post-synaptic (Dh44) neurons likely drive sleep-wake rhythm circuit development. Our CCHa1 peptide data suggest that Dh44 neurons may undergo changes in CCHa1-R expression or subcellular localization between the L2 and L3 stages: we only observed an increase in Dh44 neural activity in response to the bath application of CCHa1 peptide at the L3 stage (Figures 4E and 4F). Interestingly, this increase in activity is absent in low nutrient conditions (Figure 4G) suggesting that the larval nutritional environment may also modulate CCHa1-R localization or expression. Indeed, we observed a disruption of sleep rhythms in L3 when glucose metabolic genes are knocked down in Dh44 neurons, demonstrating that post-synaptic processes likely initiate onset of circadian sleep. These findings raise intriguing questions for how changes in an organism’s energetic and nutritional state influence sleep-circadian circuit development. Perhaps larval Dh44 neurons respond to an increase in glucose levels in the environment by promoting CCHa1-R localization to the membrane. In this model, changes in CCHa1-R subcellular localization allow Dh44 neurons to become competent to receive clock-driven cues, while this or other Dh44-derived signals promote circuit connectivity with DN1as to drive consolidation of sleep at the L3 stage. There are no available antibodies or endogenous fluorescent reporters of CCHa1-R, limiting our ability to examine receptor subcellular localization. Additionally, while our study focuses on presumed CCHa1 synaptic signaling between DN1a and Dh44 neurons, we cannot rule out the possibility of CCHa1 volume transmission from DN1as or other sources as contributors to sleep-wake regulation. Regardless, our data open avenues for future work on the molecular and subcellular mechanisms regulating DN1a-Dh44 circuit development.

Our findings demonstrate that larvae exhibit both sleep-wake and feeding daily rhythms at the L3 stage, but not earlier^24^. This raises the obvious question of whether sleep and feeding are opposite sides of the same coin. While larvae cannot eat when they are sleeping, we have observed distinct effects of certain manipulations on sleep behaviors but not feeding. For example, knockdown of *Hex-C* in Dh44 neurons disrupts sleep rhythms with no obvious effect on feeding behavior (Figure 5D). It is, of course, possible to affect both sleep and feeding behaviors with the same manipulation (e.g., activation of NPF+ neurons) underscoring that they are highly inter-connected behaviors. Future work will leverage the larval system to examine how sleep-wake and feeding circuitry communicate to balance these rhythmic behaviors across developmental periods.

## Acknowledgements

We thank members of the Kayser Lab, Raizen Lab, and other members of the Penn Chronobiology and Sleep Institute for helpful discussions and input. Figure S6 was created with Biorender.com.

## Funding

This work was supported by NIH DP2NS111996, NIH R01NS120979, and a Burroughs Wellcome Career Award for Medical Scientists to M.S.K.; Hartwell Foundation Fellowship to A.R.P.

## Author contributions

Conceptualization, A.R.P., M.S.K.; Investigation, A.R.P., L.Z., E.V., S.H.T.; Writing – Original Draft, A.R.P. and M.S.K.; Writing – Review & Editing, all authors; Project supervision and funding, M.S.K.

## Data and Materials Availability

All data needed to evaluate the conclusions in the paper are present in the paper and/or the Supplementary Materials.

## Competing Interests

All authors declare that they have no competing interests.

## Materials and Methods

### ly Stocks

The following lines have been maintained as lab stocks or were obtained from Dr. Amita Sehgal: iso31, tim0^25^, Dh44^VT^-Gal4 (VT039046)^41^, cry-Gal4 pdf-Gal80^42^, UAS-TrpA1^43^, UAS-mCherry RNAi, LexAOP-GCaMP6 UAS-P2X2^34^, and UAS-GCaMP7f. Dh44-LexA (80703), npf-Gal4 (25681), R76G11-Gal4 (48333), Hex-C RNAi (57404), Glut1 RNAi (40904), PyK RNAi (35218), GCN2 RNAi (67215), and crc RNAi (80388) were from the Bloomington *Drosophila* Stock Center (BDSC).

### Larval rearing and sleep assays

Larval sleep experiments were performed as described previously^24,44^. Briefly, molting 2^nd^ instar or 3^rd^ instar larvae were placed into individual wells of the LarvaLodge containing either 120 µl (for L2) or 95 µl (for L3) of 3% agar and 2% sucrose media covered with a thin layer of yeast paste. The LarvaLodge was covered with a transparent acrylic sheet and placed into a DigiTherm (Tritech Research) incubator at 25°C for imaging. Experiments were performed in the dark. For thermogenetic experiments, adult flies were maintained at 22°C. Larvae were then placed into the LarvaLodge (as described above) which was moved into a DigiTherm (Tritech Research) incubator at 30°C for imaging.

### LarvaLodge image acquisition and processing

Images were acquired every 6 seconds with an Imaging Source DMK 23GP031 camera (2592 × 1944 pixels, The Imaging Source, USA) equipped with a Fujinon lens (HF12.55A-1, 1:1.4/12.5 mm, Fujifilm Corp., Japan) with a Hoya 49mm R72 Infrared Filter as described previously^24,44^. We used IC Capture (The Imaging Source) to acquire time-lapse images. All experiments were carried out in the dark using infrared LED strips (Ledlightsworld LTD, 850 nm wavelength) positioned below the LarvaLodge.

Images were analyzed using custom-written MATLAB software (see Churgin et al 2019^45^ and Szuperak et al 2018^44^). Temporally adjacent images were subtracted to generate maps of pixel value intensity change. A binary threshold was set such that individual pixel intensity changes that fell below 40 gray-scale units within each well were set equal to zero (“no change”) to eliminate noise. For 3^rd^ instars, the threshold was set to 45 to account for larger body size. Pixel changes greater than or equal to threshold value were set equal to one (“change”). Activity was then calculated by taking the sum of all pixels changed between images. Sleep was defined as an activity value of zero between frames. For 2^nd^ instar sleep experiments done across the day, total sleep was summed over 6 hrs beginning 2 hrs after the molt to second instar. For sleep experiments performed at certain circadian times, total sleep in the 2^nd^ hour after the molt to second (or third) instar was summed.

### Feeding behavior analysis

For feeding rate analysis, newly molted 2^nd^ instar or 3^rd^ instar larvae were placed in individual wells of the LarvaLodge containing 120 µl of 3% agar and 2% sucrose media covered with a thin layer of yeast paste. Larvae were then imaged continuously with a Sony HDR-CX405 HD Handycam camera (B&H Photo, Cat. No: SOHDRCX405) for 5 minutes. The number of mouth hook contractions (feeding rate) was counted manually over the imaging period and raw numbers were recorded. For food intake analysis, newly molted 2^nd^ instar or 3^rd^ instar larvae were starved for 1 hr in petri dishes with water placed on a Kimwipe. To compare groups of larvae of similar body weights, 13 L3 larvae and 26 L2 larvae were grouped together. Larvae were placed in a petri dish containing blue-dyed 3% agar, 2% sucrose, and 2.5% apple juice with blue-dyed yeast paste on top for 4 hrs at 25°C in constant darkness. We found that 4 hrs on blue-dyed agar was sufficient to reflect total food consumption in each condition with a shorter period of time (1 hour) causing more variability. After 4 hrs, groups of larvae were washed in water, put in microtubes, and frozen at -80°C for 1 hr. Frozen larvae were then homogenized in 300 µl of distilled water and spun down for 5 min at 13,0000 rpm. The amount of blue dye in the supernatant was then measured using a spectrophotometer (OD_629_). Food intake represents the OD value of each measurement.

### Aversive Olfactory conditioning

We used an established two odor reciprocal olfactory conditioning paradigm with 10 mM quinine (quinine hydrochloride, EMSCO/Fisher, Cat. No: 18-613-007) as a negative reinforcement to test short-term or long-term memory performance in L2 and early L3 larvae^46^ at CT12-15^24^. Experiments were conducted on assay plates (100 × 15 mm, Genesee Scientific, Cat. No: 32-107) filled with a thin layer of 2.5% agarose containing either pure agarose (EMSCO/Fisher, Cat. No: 16500-500) or agarose plus reinforcer. As olfactory stimuli, we used 10 µl amyl acetate (AM, Sigma-Aldrich, Cat. No: STBF2370V, diluted 1:50 in paraffin oil-Sigma-Aldrich, Cat. No: SZBF140V) and octanol (OCT, Fisher Scientific, Cat. No: SALP564726, undiluted). Odorants were loaded into the caps of 0.6 mL tubes (EMSCO/Fisher, Cat. No: 05-408-123) and covered with parafilm (EMSCO/Fisher, Cat. No: 1337412). For naïve preferences of odorants, a single odorant was placed on one side of an agarose plate with no odorant on the other side. A group of 30 larvae were placed in the middle. After 5 minutes, individuals were counted on the odorant side, the non-odorant side, or in the middle. The naïve preference was calculated by subtracting the number of larvae on the non-odorant side from the number of larvae on the odorant side and then dividing by the total number of larvae. For naïve preference of quinine, a group of 30 larvae were placed in the middle of a half agarose-half quinine plate. After 5 minutes, individuals were counted on the quinine side, the agarose side, or in the middle. The naïve preference for quinine was calculated by subtracting the number of larvae on the quinine side from the number of larvae on the agarose side and then dividing by the total number of larvae. Larvae were trained by exposing a group of 30 larvae to AM while crawling on agarose medium plus quinine reinforcer. After 5 min, larvae were transferred to a fresh Petri dish containing agarose alone with OCT as an odorant (AM+/OCT). A second group of 30 larvae received the reciprocal training (AM/OCT+). Three training cycles were used for all experiments. For long-term memory, larvae were transferred after training onto agarose plates with a small piece of Kimwipe moistened with tap water and covered in dry active yeast (LabScientific, Cat. No: FLY804020F). Larvae were then kept in the dark for 1.5 hrs before testing memory performance. Training and retention for thermogenetic experiments were conducted at 30°C. For short-term memory, larvae were immediately transferred after training onto test plates (agarose plus reinforcer) on which AM and OCT were presented on opposite sides of the plate. After 5 min, individuals were counted on the AM side, the OCT side, or in the middle. We then calculated a preference index (PREF) for each training group by subtracting the number of larvae on the conditioned stimulus side from the number of larvae on the unconditioned stimulus side. For one set of experiments, we calculated two PREF values: 1a) PREF_AM+/OCT_ = (#AM - #OCT)/ # TOTAL; 1b) PREF_AM/OCT+_ = (#OCT-#AM)/ # TOTAL. We then took the average of each PREF value to calculate an associative performance index (PI) as a measure of associative learning. PI = (PREF_AM+/OCT_ + PREF_AM/OCT+)_/2.

### Arousal threshold

Blue light stimulation was delivered as described in ^24,44^ using 2 high power LEDs (Luminus Phatlight PT-121, 460 nm peak wavelength, Sunnyvale, CA) secured to an aluminum heat sink. The LEDs were driven at a current of 0.1 A (low intensity). We used a low intensity stimulus for 4 sec every 2 minutes for 1 hr beginning the 2^nd^ hr after the molt to second (or third) instar. We then counted the number of larvae that showed an activity change in response to stimulus. The percentage of animals that moved in response to the stimulus was recorded for each experiment. For each genotype, at least 4 biological replicates were performed. We then plotted the average percentage across all replicates.

### P2X2 Activation and GCaMP imaging

All live imaging experiments (P2X2 and CCHa1 bath application) were performed as described previously^24^. Briefly, brains were dissected in artificial hemolymph (AHL) buffer consisting of (in mM): 108 NaCl, 5 KCl, 2 CaCl2, 8.2 MgCl2, 4 NaHCO3, 1 NaH2PO4-H20, 5 Trehalose, 10 Sucrose, 5 HEPES, pH=7.5. Brains were placed on a small glass coverslip (Carolina Cover Glasses, Circles, 12 mm, Cat. No: 633029) in a perfusion chamber filled with AHL.

For P2X2 imaging, dissections were performed at CT12-15 and AHL buffer was perfused over the brains for 1 min of baseline GCaMP6 imaging, then ATP was delivered to the chamber by switching the perfusion flow from the channel containing AHL to the channel containing 2.5 mM ATP in AHL, pH 7.5. ATP was perfused for 2 min and then AHL was perfused for 2 min. Twelve-bit images were acquired with a 40 × water immersion objective at 256 × 256-pixel resolution. Z-stacks were acquired every 5 sec for 3 min. Image processing and measurement of fluorescence intensity was performed in ImageJ as described previously^24^. For each cell body, fluorescence traces over time were normalized using this equation: ΔF/F = (F_n_-F_0_)/F_0_, where F_n_=fluorescence intensity recorded at time point n, and F_0_ is the average fluorescence value during the 1 min baseline recording. Maximum GCaMP change (ΔF/F) for individual cells was calculated using this equation: ΔF/F_max_ = (F_max_-F_0_)/F_0_, where F_max_=maximum fluorescence intensity value recorded during ATP application, and F_0_ is the average fluorescence value during the 1 min baseline recording. All analysis was done blind to experimental condition.

For CCHa1 bath application, dissections were performed at CT12-15 and AHL buffer was perfused over the brains for 1 min of baseline GCaMP7f imaging, then CCHa1 peptide was delivered to the chamber by switching the perfusion flow from the channel containing AHL to the channel containing 1 µM synthetic CCHa1 in AHL, pH 7.5. CCHa1 was perfused for 2 min, followed by a 1 min wash-out with AHL. For the AHL negative control, the perfusion flow was switched from one channel containing AHL to another channel containing AHL. Twelve-bit images were acquired with a 40 × water immersion objective at 256 × 256-pixel resolution. Z-stacks were acquired every 10 sec for 4 min. Image processing and measurement of fluorescence intensity was performed in ImageJ. A max intensity Z-projection of each time step and Smooth thresholding was used for analysis. Image analysis was performed in a similar manner as for the P2X2 experiments. All analysis was done blind to experimental condition.

### Gaboxadol treatment

Early second or third instar larvae were starved for 1 hour and then fed 75 µl of 25 mg/mL Gaboxadol (hydrochloride) (Thomas Scientific, Cat No: C817P41) in diluted yeast solution for 1 hour prior to loading in LarvaLodge containing 120 µl of 3% agar and 2% sucrose media covered with a thin layer of 25 mg/mL Gaboxadol yeast paste. For LTM experiments, starved early second instars were fed 25 mg/mL Gaboxadol for 1 hour prior to training and maintained on 25 mg/mL Gaboxadol in diluted yeast solution during retention period.

### Dietary Manipulations

Fly food was prepared using the following recipes (based on Poe et al 2020)^47^:

| Ingredients | Control food (444 mM glucose) | Low sugar (L.S.) (66 mM glucose) |
| --- | --- | --- |
| Distilled H<sub>2</sub>O | 234 mL | 234 mL |
| Agar | 2 g (10g/L) | 2 g (10g/L) |
| Glucose | 20 g | 3 g |
| Inactive yeast | 20 g | 20 g |
| Acid mix (phosphoric acid<br>+ propionic acid) | 2 mL | 2 mL |
| Target final solution<br>volume | 250 mL | 250 mL |

Acid Mix was made by preparing Solution A (41.5 ml Phosphoric Acid mixed with 458.5 ml distilled water) and Solution B (418 ml Propionic Acid mixed with 82 ml distilled water) separately and then mixing Solution A and Solution B together.

Adult flies were placed in an embryo collection cage (Genesee Scientific, cat#: 59-100) and eggs were laid on a petri dish containing either control (ctrl) or Low sugar (L.S.) food. Animals developed on this media for three days.

### Larval Body Weight and Length Measurements

For weight, groups of 5 early 3^rd^ instar larvae raised on either control- or low sugar (L.S.)-filled petri dishes were washed in tap water and dried using a Kimwipe. The 5 larvae were then weighed as a group on a scale and the weight in mg was recorded. For the Gaboxadol experiments, groups of 10 early 2^nd^ instar larvae or groups of 5 early 3^rd^ instar larvae were weighed. For length, images of individual early 3^rd^ instar larvae in the LarvaLodge were measured in ImageJ (Fiji) using the straight line tool. The total body length was determined in pixels for individual larvae on each condition.

### Statistical analysis

All statistical analysis was done in GraphPad (Prism). For comparisons between 2 conditions, two-tailed unpaired *t*-tests were used. For comparisons between multiple groups, ordinary one-way ANOVAs followed by Tukey’s multiple comparison tests were used. For comparisons between different groups in the same analysis, ordinary one-way ANOVAs followed by Sidak’s multiple comparisons tests were used. For comparisons between time and genotype, two-way ANOVAs followed by Sidak’s multiple comparisons tests were used. For comparison of GCaMP signal in CCHa1 experiments, Mann-Whitney test was used. \**P*<0.05, \*\**P*<0.01, \*\*\**P*<0.001. Representative confocal images are shown from at least 8-10 independent samples examined in each case.

---

**Supplemental Figure 1:**
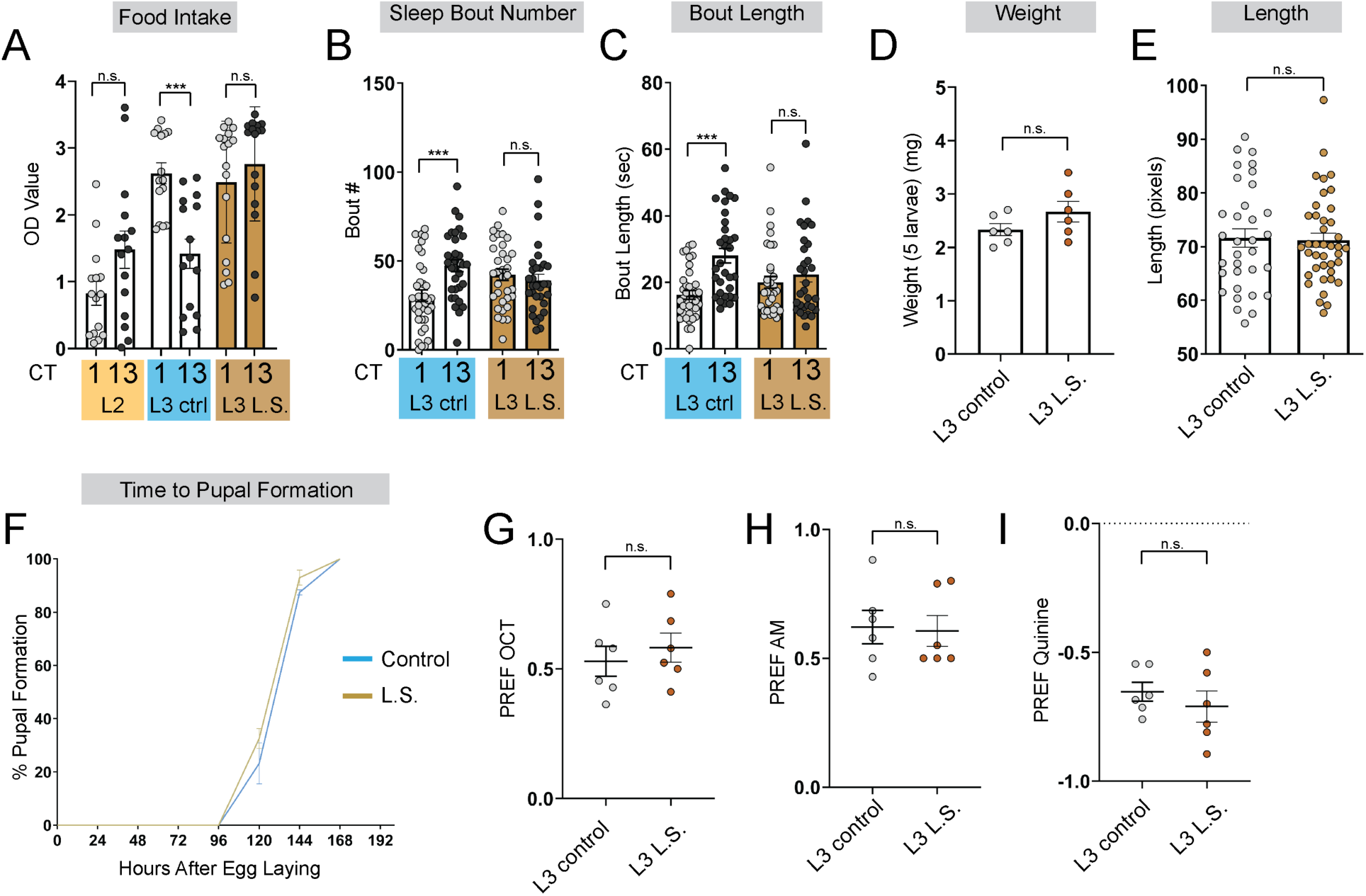
Larvae reared on low sugar diet develop normally. **(A)** Feeding amount of L2 controls, L3 raised on regular (ctrl) food, and L3 raised on low sugar (L.S.) food at CT1 and CT13. **(B, C)** Sleep bout number (B) and bout length (C) at CT1 and CT13 in L3 raised on regular (ctrl) and L.S. food. **(D)** Total body weight of early L3 (in groups of 5) raised on ctrl or L.S. food. **(E)** Total body length of early L3 raised on ctrl or L.S. food. **(F)** Developmental analysis of time to pupal formation of animals raised on ctrl or L.S. food. **(G-I)** Naïve OCT, AM, and quinine preference in L3 raised on ctrl and L.S. food. B-C, n=29-34 larvae; D, n=30 larvae per food condition; E, n=33-40 larvae; F, n=100-170 larvae; G-I, n=6 PREFs (180 larvae) per genotype. Two-way ANOVAs followed by Sidak’s multiple comparison test [(B-C)]; Unpaired two-tailed Student’s *t*-tests [(D-E) and (G-I)].

**Supplemental Figure 2:**
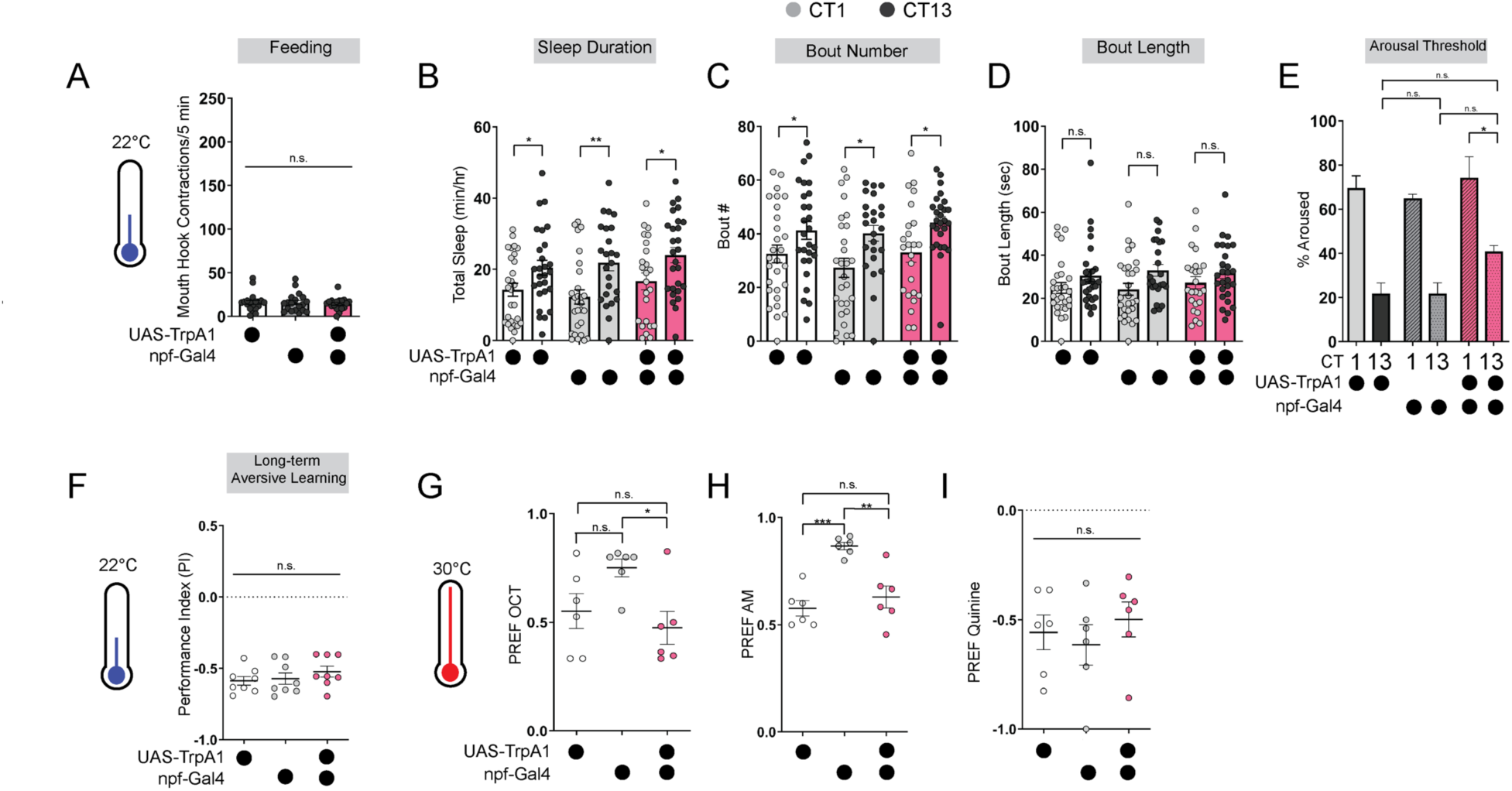
Baseline odor preferences, feeding, and sleep are not affected by *npf*-Gal4 manipulations. **(A)** Feeding rate of L3 expressing *npf-*Gal4>*UAS-TrpA1* and genetic controls at 22°C at CT13. **(B-E)** Sleep duration (B), bout number (C), bout length (D), and arousal threshold (E) in L3 expressing *npf-*Gal4>*UAS-TrpA1* and genetic controls at CT1 and CT13 at 22°C (temperature controls). **(F)** Long-term aversive memory performance in L3 expressing *npf-*Gal4>*UAS-TrpA1* and genetic controls at 22°C (temperature controls). **(G-I)** Naïve OCT, AM, and quinine preference in L3 expressing *npf-*Gal4>*UAS-TrpA1* and genetic controls at 30°C. A, n=18-20 larvae; B-D, n=22-27 larvae; E, n=120-205 sleep episodes, 18 larvae per genotype; F, n=8 PIs (240 larvae) per genotype; G-I, n=6 PREFs (180 larvae) per genotype. One-way ANOVAs followed by Sidak’s multiple comparisons tests [(A), (E), and (F-I)]; Two-way ANOVAs followed by Sidak’s multiple comparison test [(B-D)].

**Supplemental Figure 3:**
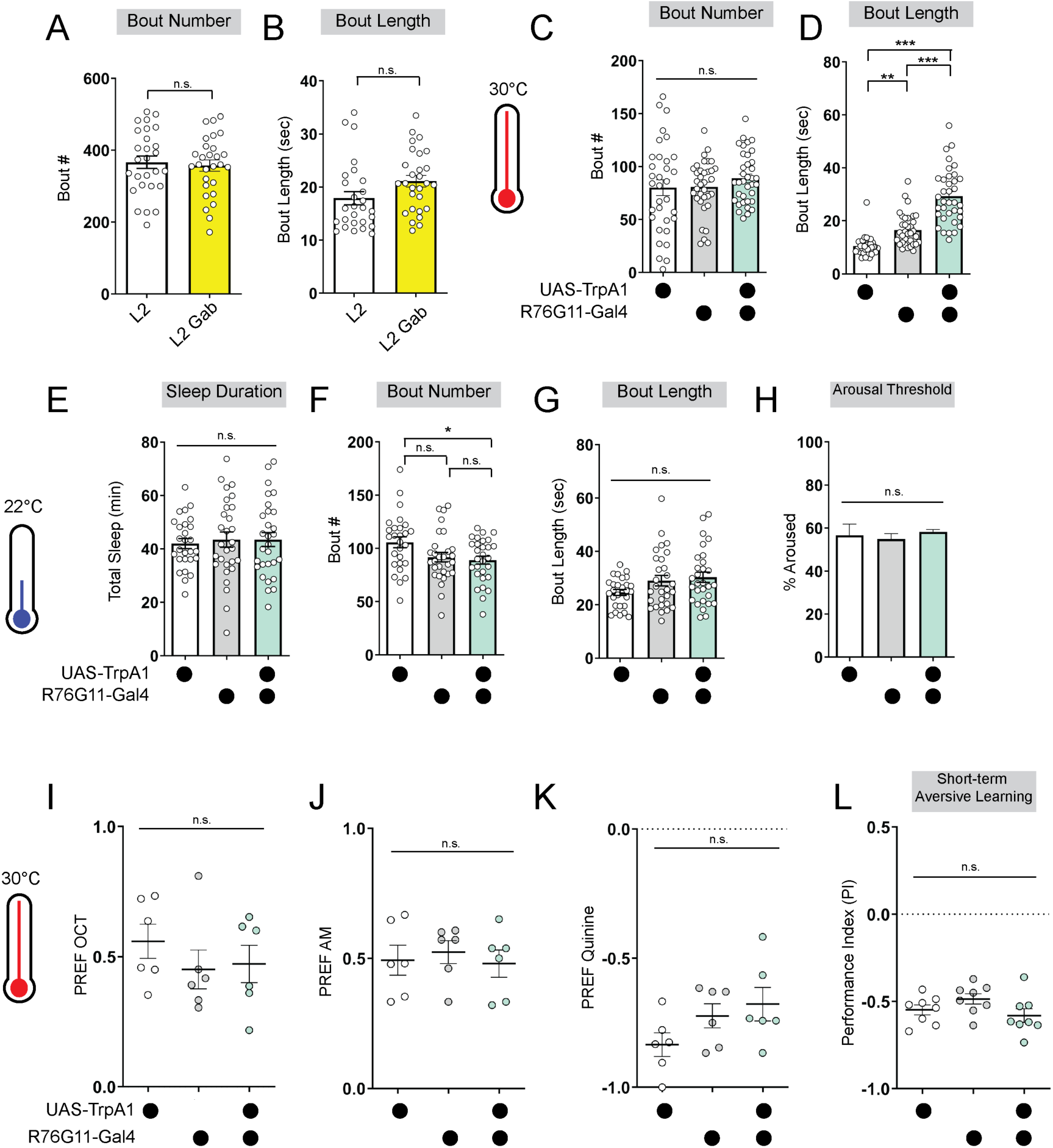
Baseline sleep and odor preferences are not disrupted by *R76G11*-Gal4 manipulations. **(A, B)** Sleep bout number (A) and bout length (B) of L2 control fed vehicle control (L2) or Gaboxadol (L2 Gab). **(C, D)** Sleep bout number (C) and bout length (D) of L2 expressing *R76G11-*Gal4>*UAS-TrpA1* and genetic controls at 30°C. **(E-H)** Sleep duration (E), bout number (F), bout length (G), and arousal threshold (H) of L2 expressing *R76G11-*Gal4>*UAS-TrpA1* and genetic controls at 22°C (temperature controls). **(I-K)** Naïve OCT, AM, and quinine preference in L2 expressing *R76G11-* Gal4>*UAS-TrpA1* and genetic controls at 30°C. **(L)** Short-term aversive memory performance of L2 expressing *R76G11-*Gal4>*UAS-TrpA1* and genetic controls at 30°C. A, B, n=28 larvae; C-D, n=33-36 larvae; E-G, n=27-29 larvae; H, n=145-316 sleep episodes, 30-40 larvae per genotype; I-K, n=6 PREFs (180 larvae) per genotype; L, n=8 PIs (240 larvae) per genotype. Unpaired two-tailed Student’s *t*-tests [(A-B)]; One-way ANOVAs followed by Sidak’s multiple comparisons tests [(C-L)].

**Supplemental Figure 4:**
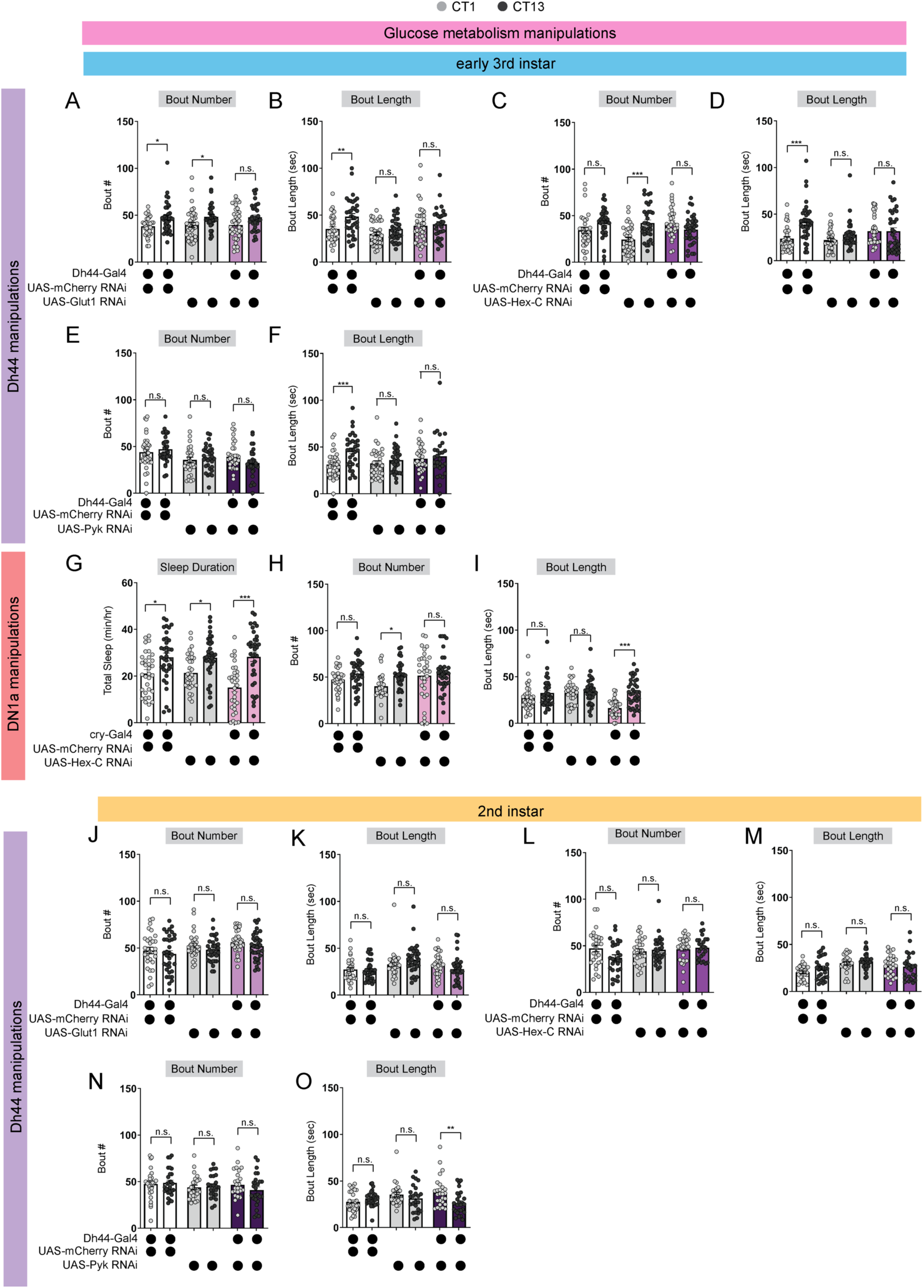
Glucose metabolic gene manipulations affect L3 sleep. **(A, B)** Sleep bout number (A) and bout length (B) in L3 expressing *UAS-Glut1-RNAi* with *Dh44-Gal4* and genetic controls at CT1 and CT13. **(C, D)** Sleep bout number (C) and bout length (D) in L3 expressing *UAS-Hex-C-RNAi* with *Dh44-Gal4* and genetic controls at CT1 and CT13. **(E, F)** Sleep bout number (E) and bout length (F) in L3 expressing *UAS-PyK-RNAi* with *Dh44-Gal4* and genetic controls as CT1 and CT13. **(G-I)** Sleep duration (G), sleep bout number (H), and bout length (I) in L3 expressing *UAS-Hex-C-RNAi* with *cry-Gal4* and genetic controls at CT1 and CT13. **(J, K)** Sleep bout number (J) and bout length (K) in L2 expressing *UAS-Glut1-RNAi* with *Dh44-Gal4* and genetic controls at CT1 and CT13. **(L, M)** Sleep bout number (L) and bout length (M) in L2 expressing *UAS-Hex-C-RNAi* with *Dh44-Gal4* and genetic controls at CT1 and CT13. **(N, O)** Sleep bout number (N) and bout length (O) in L2 expressing *UAS-PyK-RNAi* with *Dh44-Gal4* and genetic controls at CT1 and CT13. A-O, n=32-40 larvae. Two-way ANOVAs followed by Sidak’s multiple comparison test [(A-O)].

**Supplemental Figure 5:**
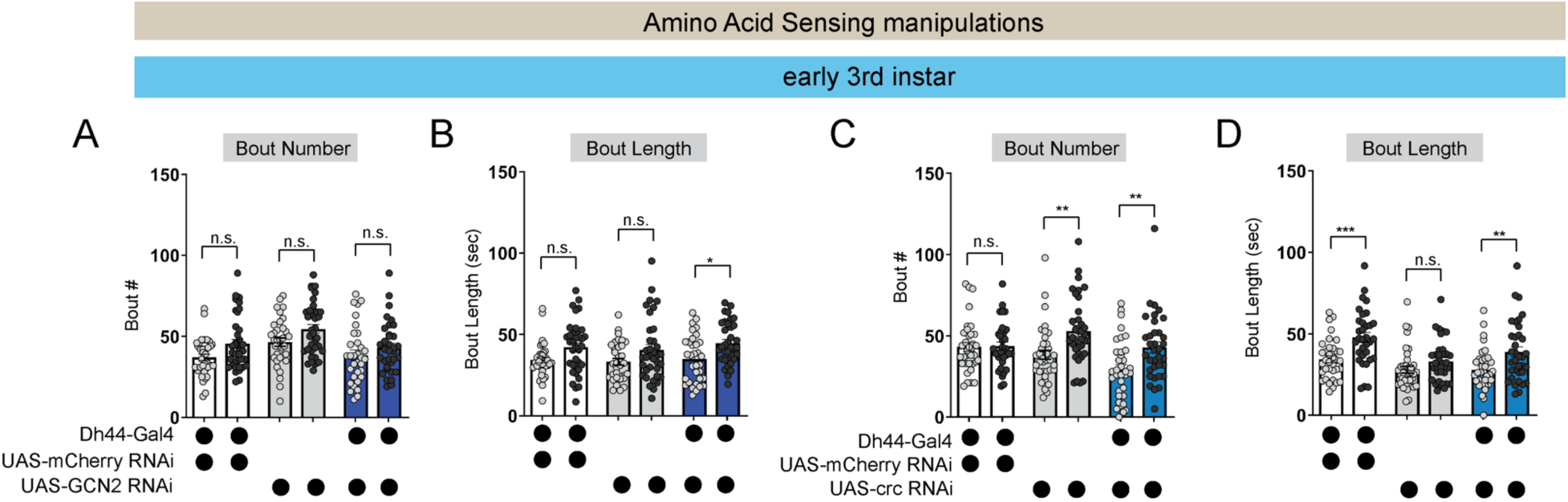
Amino acid sensing gene manipulations do not affect L3 sleep. **(A, B)** Sleep bout number (A) and bout length (B) in L3 expressing *UAS-GCN2-RNAi* with *Dh44-Gal4* and genetic controls at CT1 and CT13. **(C, D)** Sleep bout number (C) and bout length (D) in L3 expressing *UAS-crc-RNAi* with *Dh44-Gal4* and genetic controls at CT1 and CT13. A-D, n=32-40 larvae. Two-way ANOVAs followed by Sidak’s multiple comparison test [(A-D)].

**Supplemental Figure 6:**
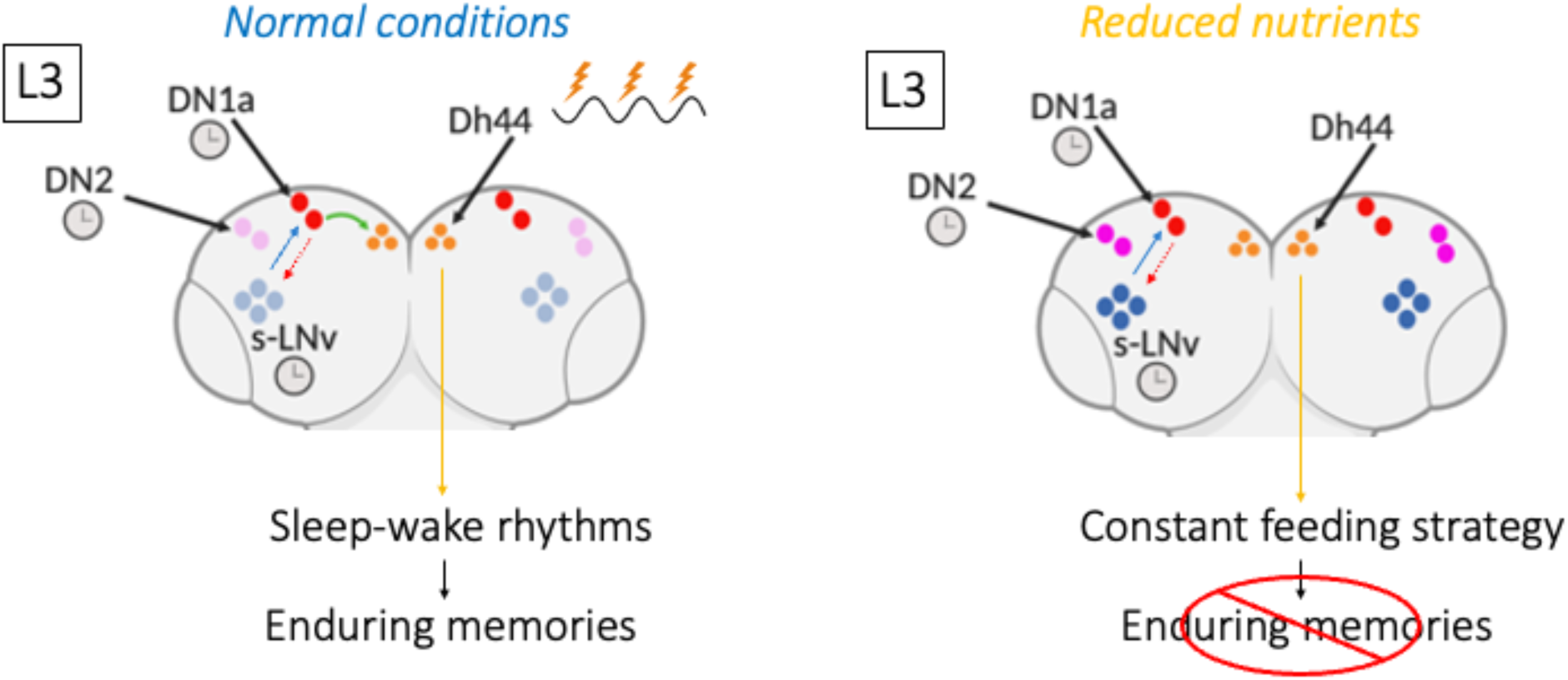
Model Figure. Clock cells in the larval brain (s-LNv, DN2, and DN1a) communicate to coordinate circadian rhythms. In third instar larvae (L3), a new connection is formed between DN1as and Dh44 neurons, generating daily neural activity rhythms in Dh44 cells that drive sleep-wake patterns, deep sleep, and more enduring memories. In the setting of reduced nutrient availability, the functional connection between the clock (DN1a) and arousal output (Dh44) is not present, facilitating a more constant feeding strategy that benefits the animal under such conditions. However, without clock control of sleep at this stage, deep sleep is lost, as is the ability to exhibit long-term memory.

## Notes

### Competing Interest Statement

The authors have declared no competing interest.

### Summary of Updates

Changes to text of manuscript and additional data

